# Identification of the sex determination region and the development of a marker to distinguish males and females in megai abalone (*Haliotis gigantea*)

**DOI:** 10.1101/2024.08.29.610404

**Authors:** Taito Kina, Motoyuki Hara, Shotaro Hirase, Kiyoshi Kikuchi

## Abstract

Sex identification markers are of considerable value to aquaculture and fisheries as they enable pre-maturity prediction of the genotypic sex of individuals as well as of the sex ratio in offspring. For these reasons, extensive investigations of the mechanisms of sex determination have been conducted in commercially important finfish species, resulting in the development of sex identification markers for many of these. However, research on shellfish is still in its infancy stage. In this study, we investigated the genomic basis of sex determination in megai abalone (*Haliotis gigantea*), a gastropod species of economic importance in fisheries and aquaculture in East Asia, with the aim of developing a sex identification marker. Initially, we examined the applicability of sex identification markers reported for two other *Haliotis* species, but found they were not transferable to *H. gigantea.* We then analyzed the sex chromosomes in *H. gigantea* using a whole-genome resequencing approach. By calculating the fixation index value between 10 females and 10 males, we identified a genomic region spanning approximately 2 Mb that exhibited nucleotide sequence divergence between the sexes (from the position 260,372 bp to 1,930,278 bp in chromosome 18). This region was located next to the pseudoautosomal region which is characterized by an absence of sequence differentiation between the sexes. In the divergent region, male-specific SNPs were identified but no female-specific SNPs were detected, indicating that this species has an XX-XY sex determination system. By exploring Indel polymorphisms in the divergent region, we successfully developed a codominant PCR marker for sex identification that could be applied to wild individuals from geographically distant locations. A cross-species analysis suggested that the sex determination region in *H. gigantea* was unlikely to be shared with the closely related species *H. discus hannai*. This raises the possibility of a widespread, rapid turnover of sex chromosomes or sex determination regions in abalones, similar to certain vertebrate lineages such as finfish.

## 1. Introduction

Western Pacific abalones are distributed along the coasts of the Japanese archipelago and the Korean Peninsula and form a species complex that includes four species or subspecies: *Haliotis discus hannai* (Ezo abalone), *H. discus discus* (kuro abalone), *H. madaka* (madaka abalone), and *H. gigantea* (megai abalone) (Ino, 1952). These species are economically important in fisheries and aquaculture in East Asia (Arai and Okumura., 2013; Cook, 2019; Gao et al., 2023: Park and Kim, 2013).

In recent years, there has been a significant decline in abalone catches in Japan and for other abalone species around the world. For instance, abalone catches in Japan decreased from approximately 6,500 tons in 1970 to around 1,200 tons in 2012 (Yamazaki and Kamoshida, 2018). To mitigate this problem, the Japanese government introduced a marine stock enhancement program in the 1970s (Hara, 2008a; Kitada 2020: Masuda and Tsukamoto, 1998), and efficient propagation techniques for abalones were established (e.g., Kikuchi and Uki, 1974; Morse et al., 1977; Uki and Kikuchi, 1984).

In addition to the stock enhancement program, attempts have been made to improve aquaculture of the species and to develop genetically improved strains (Hara, 1992; Li et al., 2018; Kawahara et al., 1999; Park and Kim, 2013; Gao et al., 2023). A large aquaculture industry has now been developed in some eastern Asian regions such as China and South Korea (Cook, 2023; Gao et al., 2023; Park and Kim, 2013; You, 2023). The recent listing of western Pacific abalone species as endangered by the International Union for Conservation of Nature and Natural Resources (IUCN) (Peters & Rogers-Bennett; 2021, Peters et al., 2022a, b, c) has further stimulated the importance for the development of genetically improved strains for aquaculture.

Much of the recent research on the species complex of western Pacific abalones has focused on *H. discus hannai* (e.g., Chen et al., 2017; Nam et al., 2017; Yu et al., 2018). However, this species is found in the northern parts of the Japanese archipelago, while the other three species/subspecies are distributed in more southern locations (Hara and Sekino, 2005; Ino, 1952; Nam et al., 2021); thus, the potential of the other species/subspecies for aquaculture in warmer climates needs to be reevaluated (Ahmed et al., 2013; Koike et al., 1988; Lyu et al., 2021). This need is especially important in an era of global warming. Among the southern species, *H. gigantea* is potentially important for temperate aquaculture as the growth of *H. gigantea* is better than that of *H. discus discus* under warmer temperature (Komazawa et al., 2004). *H. madaka* is a rare species and there is little information of relevance to breeding experiments.

In aquaculture, the ability to control sex and reproduction is crucial for efficient propagation of the target species, as these factors influence growth and product quality, as well as quantity (Budd et al., 2015; Kikuchi and Uki: 1974, Martínez et al., 2014; Piferrer et al., 2012). Notable advances have been made in recent years on genetic sex identification (Li et al., 2022). For example, more than ten master sex-determining genes have been identified in economically important fish in the last 20 years (Kikuchi and Hamaguchi, 2013; Li et al., 2022; Nagahama et al., 2021). These results have revealed a remarkable diversity in the genetic mechanisms of sex determination in fish, contrasting with the largely conserved process found in mammals (Graves and Peichel, 2010; Nagahama et al., 2021). The identification of these genes not only enhanced our understanding of the mechanisms of fish sex determination, but also led to the development of sex identification markers, which are now being exploited for controlled breeding, selective breeding, the development of monosex stocks, and the management of wild stocks in aquaculture and fisheries (Akita et al., 2020; Budd et al., 2015; Ieda et al., 2018; Li et al., 2022; Moore et al., 2020; Rondeau et al., 2013).

In the genus *Haliotis*, which includes more than 50 species (Geiger, 2000), genetic markers capable of identifying males and females have been reported for only *H. discus hannai* (Luo et al., 2021; Yang et al., 2019) and *H. diversicolor* (Weng et al., 2022). *H. discus hannai* is estimated to have diverged from *H. gigantea* approximately 14 million years ago (Hirase et al., 2021), while *H. diversicolor* diverged much earlier (Lee and Vaequier, 1995; Streit et al., 2006). Therefore, it is more likely that the markers identified in *H. discus hannai* could be used for *H. gigantea* than those of *H. diversicolor*. However, the transferability of the markers from both species has yet to be assessed. Moreover, the sex-determining system or sex chromosome(s) in *H. gigantea* have not yet been analyzed.

Sexual dimorphism for growth has not been reported in *H. gigantea*; thus, there is no need for monosex populations in large-scale aquaculture production. However, for genetic improvement programs, where controlled crosses are necessary and the assessment of breeding values is preferably performed at the juvenile stage (e.g., Hosoya et al., 2021), early sexing of individuals using genetic markers will be valuable. This is particularly true given that sex identification in this species is only possible after they have been in culture for three years (Mie Prefectural Fisheries Experiment Station, 2014; Yamakawa et al., 1994). In addition, to avoid genetic risks involved in the use of sex-reversed individuals (e.g., males with XX genotypes) in stock enhancement programs, a reliable method for identifying the genotypic sex is required (Ieda et al., 2018; Kanaiwa et al., 2002; Stelkens and Wedekind, 2010).

In this study, we initially attempted to use the reported sex identification markers for *H discus hannai* and *H. diversicolor* in *H. gigantea*. The markers were not found to be transferable to *H. gigantea.* We subsequently analyzed the sex chromosomes in *H. gigantea* using a whole-genome resequencing approach and successfully developed a codominant genetic marker for sex identification.

## 2. Materials and Methods

### 2.1. Samples

The megai abalone (*H. gigantea*) used in this study were collected from the coast of Mie prefecture (n = 15 for each sex) and Tokushima prefecture (n = 5 for each sex), Japan (Table S1). Ezo abalone (*H. discus hannai*) (n = 5 for each sex) were collected from the coast of Hokkaido, Japan. Small abalone (*H. diversicolor*) (n = 5 for each sex) were collected from the coast of Oita prefecture, Japan. The collected specimens were promptly dissected, and their phenotypic sex was tentatively determined through visual observation of the gonads (Figure 1). The gonads were then removed and fixed in Davidson’s fixative at room temperature. The tissues were sectioned at a thickness of 10 µm and the sections were stained with hematoxylin and eosin for histological analysis to confirm the phenotypic sex. To obtain DNA, foot tissues were excised, cut into 1 cm cubes, and then kept at room temperature in TNU8E buffer (10 mM Tris–HCl, pH 7.5; 125 mM NaCl; 10 mM EDTA; 1% SDS; 8 M urea) (Asahida et al., 1996) until required.

**Figure 1.**
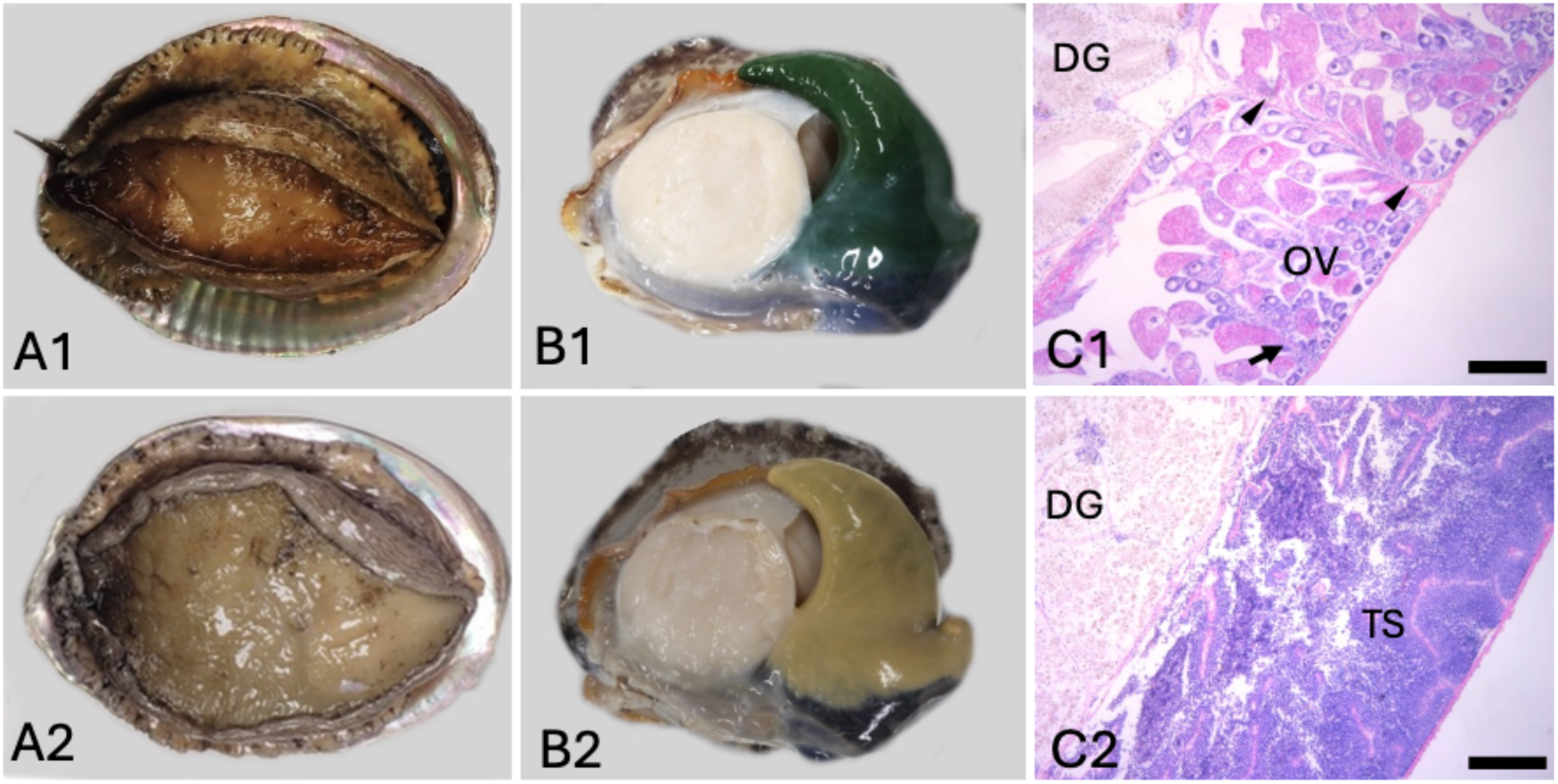
The ovary and testis of megai abalone (*Haliotis gigantea*). (A1) The ventral surfaces of a female individual and of a male individual (A2). (B1) The dorsal surface of a female individual with the shell removed, exhibiting a green gonad; (B2) the dorsal surface of a male individual with a cream-colored gonad. (C1) Histological observation of the ovary; (C2) Histological section of the testis. DG, digestive gland; OV, ovary; TS, testis. Arrow head: Trabeculae extending from the ovarian wall to the wall of the digestive gland. Arrow: Trabeculae that do not reach the digestive gland wall. Scale bar = 20µm.

### 2.2. DNA extraction

Foot tissue was gently dissolved by addition of 1 µL of 50 µg/mL Proteinase K (Macherey-Nagel GmbH & Co. KG, Duren, Germany) to 1 ml TNE8U buffer containing the tissue and incubating at 55°C overnight. Using 500 µL of the solution containing the dissolved tissue, genomic DNA was extracted following the manufacturer’s protocol for the Puregene core kit A (Qiagen, Venlo, Netherlands). Subsequently, genomic DNA was purified using DNeasy PowerClean Pro (Qiagen). The concentration of the purified DNA was measured using a Qubit kit (Quantas, Promega Corporation, Madison, Wisconsin, United States).

### 2.3. PCR conditions for testing interspecific transferability of sex identification markers

The diagnostic markers developed for *H. discus hannai* (Luo et al., 2021) and *H. diversicolor* (Wang et al., 2022) were amplified by PCR using the KOD One™ PCR Master Mix (Toyobo, Osaka, Japan) with the published primers (Table S2). The reaction mixture for each sample contained 0.3 µL each of the forward and reverse primers (10 µM each), 3.4 µL of UPW, 5 µL KOD One™ PCR Master Mix, and 1 µL genomic DNA solution. The PCR conditions consisted of an initial denaturation at 94°C for 3 minutes; 34 cycles of denaturation at 98°C for 30 seconds, annealing at Tm for 30 seconds, and extension at 68°C for 30 seconds; followed by a final extension at 68°C for 1 minute. The amplified products were separated using 3% agarose gel electrophoresis.

### 2.4. Resequencing and variant calling

Genomic DNA from 20 *H. gigantea* individuals from Mie region (10 males and 10 females) was used for library preparation (Table S1) using standard procedures with the NEB Next Ultra II DNA Library Prep Kit for Illumina (New England Biolabs, Massachusetts, USA). Whole-genome sequencing was outsourced to BGI Japan (Shenzhen, Guangdong, China) and performed using DNB-seq under PE150 bp conditions, generating over 90 GB of data. The obtained read data underwent quality control using FastQC (http://www.bioinformatics.babraham.ac.uk/projects/fastqc/); paired-end data containing adapter contamination and low-quality bases were trimmed using Trimomatic v. 0.36 (Bolger et al., 2014). The clean read data were mapped to the draft genome sequence of *H. gigantea* using BWA mem (Li and Durbin, 2009). The resulting SAM files were converted to BAM files, and duplicate reads were removed using MarkDuplicates from the Picard toolkit 2.9.0 (http://broadinstitute.github.io/picard/). The aligned genome was subjected to realignment using the Genome Analysis Toolkit package (GATK) 3.7, including RealignerTargetCreator and IndelRealigner (McKenna et al., 2010). Genotyping and variant calling were performed by integrating all samples with Bcftools mpileup, and SNPs and Indels were detected. The obtained data were filtered using VCFtools with the following parameters: remove-indels, max-missing 0.5, --minDP 6, --minQ 20, -- min-alleles 2, --max-alleles 2 (Danecek et al., 2011). SNPs were also called using the ANGSD 0.939 package which is suitable for genotyping from low or medium coverage sequencing data (https://github.com/ANGSD/angsd) (-GL 2, -doGlf 2, -doDepth 1, - doCounts 1, -doMajorMinor 1, -doMaf 2, -minMaf 0.05, -uniqueOnly 1, -minMapQ 30, -minQ 20, -skipTriallelic 1, -nThreads, $ncpu, -minInd 5) (Korneliussen et al., 2014).

To assess population structure using a principal component analysis (PCA), we converted the beagle file produced from ANGSD into a cov file using PCAngsd 1.21 (https://github.com/Rosemeis/pcangsd) with the options "-sites_save, -post_save, -e 10" and generated scatter plots with the R "eigen" function and the ggloot2 package (https://github.com/tidyverse/ggplot2).

### 2.5. Identification of sex determination locus

To identify genomic regions associated with sex, we performed genome-wide association studies (GWAS) using the ANGSD 0.933 package with options of “-doAsso 2, -doPost 1, -GL 1, -doMajorMinor 1, -doMaf 1, -SNP_ pval 1e-6”. Manhattan plots were generated by the qqman in R package (https://github.com/stephenturner/qqman).

The enrichment ratio for sex-associated SNPs on each chromosome was calculated by dividing the observed number on a chromosome by the expected number. The expected number was determined using the ratio of sex-associated SNPs in the entire genome to the total number of SNPs in the entire genome. Bar graphs were created using the ggplot2 in R package. To detect sex chromosomes with genetic differentiation between males and females, we conducted *F*st analysis in 50 kb windows using ANGSD with options of “-uniqueOnly 1, -remove_bads 1, -only_proper_pairs 1, -trim 0, -C 50, -baq 1, -minMapQ 20, -minQ 20, -minInd 10, -setMinDepth 20, -setMaxDepth 200, - doCounts 1, -GL 1, -doSaf 1” and realSFS with option “fst stats2, -win 50000, -step 50000, -whichFST”. The resulting data were visualized using the qqman R package. In evolutionarily young sex chromosome systems where the Y or W is not yet extensively diverged from the X or Z, polymorphic loci are often heterozygous in one sex while homozygous in the other sex as a result of segregation according to an XX-XY or ZZ-ZW system. We searched for such SNPs (sex-specific SNPs) using a Perl script, Find_sex_loci_from_GT.pl (https://github.com/lyl8086/find_sex_loci) (Li et al., 2020) in the VCF file generated by BWA/Bcftools in section 2.3. and counted the number at a window size of 50 kb. The resulting data were visualized using the qqman R package.

### 2.6. Development of a sex identification marker

Through use of Find_sex_loci_from_GT.pl, we identified sex-specific loci in the Indel list in the VCF file generated by BWA/Bcftools and confirmed the sex-specificity in the resequencing data by visualization with the Integrative Genomic Viewer (IGV) (http://www.broadinstitute.org/software/igv/). Then the PCR primers were designed manually so as to produce the PCR amplicon spanning the targeted Indel (Table 1).

**Table 1.**
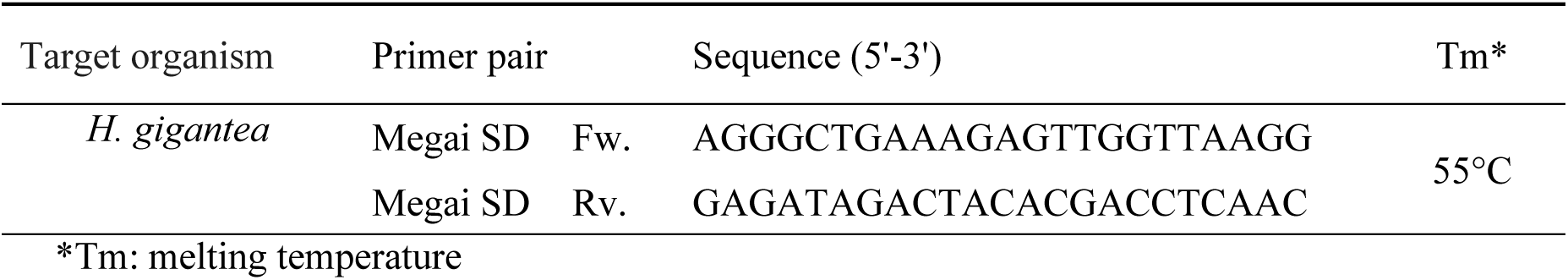
The sex-identification primer pairs based on the male-specific deletion in megai abalone (*H. gigantea*).

PCR was conducted as described in the section 2.3 except for the primers. We used *H. gigantea* specimens that were not used for resequencing (5 males and 5 females from Tokushima region; 5 males and 5 females from Mie region) in addition to the Mie samples used for resequencing analysis (5 males and 5 females)(Table S1).

## Results

### 3.1. Gonadal sex and histology

Most of the literature on gametogenesis in *H. gigantea* is focused on seasonality using oocyte diameter as a main variable (e.g., Ino, 1952; Yamakawa et al., 1994); few detailed histological studies have been reported, although Shin et al. (2020) did describe a population from a coastal area of the Korean peninsula. Our histological analysis here confirmed that the cell types and structures of the maturing ovary and testis in *H. gigantea* are similar to those reported in other abalone species (e.g., Awaji and Hamano, 2004; Giorgi et. al, 1977; Martin and Miller-Walker, 1983) (n = 5 for each sex; Figure 1). For example, follicles that contain developing oocytes associate with trabeculae that extend from the ovarian wall to the wall of the digestive gland (Figure 1 C1, arrowhead). In addition, developing oocytes are also associated with trabeculae that do not reach the digestive gland wall (Figure 1 C1, arrow) (Awaji and Hamano, 2004, Roux et al., 1983). In the maturing testis, the lumens are filled with spermatogenic cells (Figure 1 C2) (Awaji and Hamano, 2004). With regard to gross morphology, gonadal tissue coloration is consistent with histological observations reported from other abalone species: the maturing ovaries are green, and the maturing testes are cream-colored (Figure 1 B1, 2). As a result, we were able to determine the phenotypic sex of *H. gigantea* samples without ambiguity. Through the same approach, we determined the phenotypic sex of *H. discus hannai* and *H. diversicolor*.

### 3.2. Transferability of diagnostic markers from other abalone species to *H. gigantea*

Genetic markers for sex identification have been reported from *H. discus hannai* and *H*.

*diversicolor* (Luo et al., 2021; Wang et al., 2022). We performed PCR on genomic DNA from *H. gigantea* using the reported primers for these markers (Table S2) to determine whether these sequences are present in the genome of *H. gigantea*.

The two sets of primers for *H. discus hannai* produced amplicons from *H. gigantea* of 250 bp and 465 bp (Figure 2); these products were of the expected size given the high similarity of the genome sequences between these two abalone species (An et al., 2005; Hirase et al., 2021). However, no amplicon length polymorphisms were detected in gel electrophoresis, indicating that the sex-linked Indels reported for *H. discus hannai* are absent in *H. gigantea*. The two sets of primers from *H. diversicolor* produced only non-specific amplicons (SC9 in Figure 2) or no amplicons from *H. gigantea* (SD3 in Figure 2). This is likely due to sequence differences at the primer sites between these two distantly related species, which is further considerd in the discussion section with Table S5 (Estes, 2005; Geiger, 2000; Streit et al., 2006).

**Figure 2.**
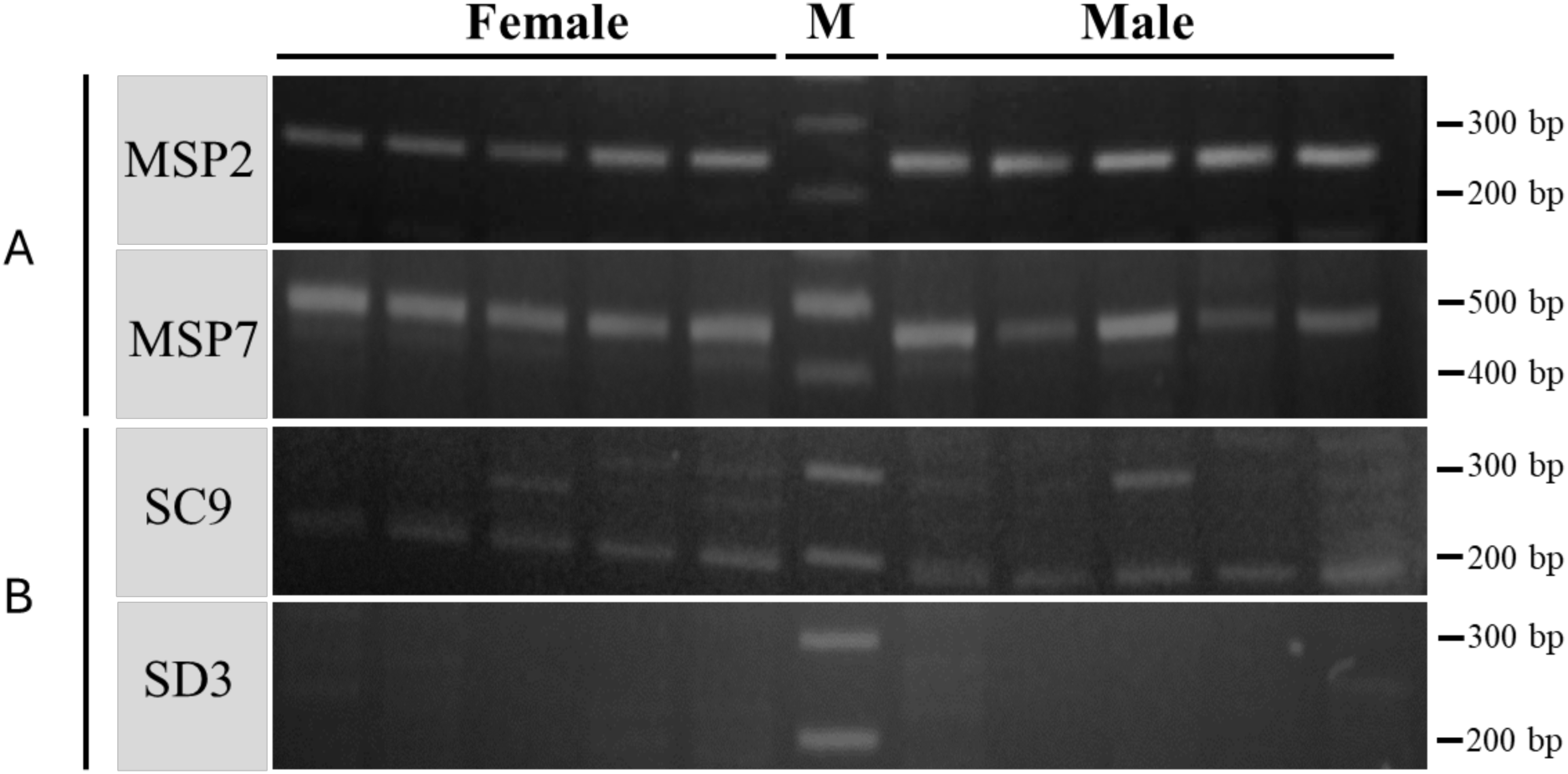
PCR products from megai abalone (*H. gigantea*) with the sex-identification primers for Ezo abalone (*H. discus hannai*) and small abalone *(H. diversicolor*). *H. gigantea* samples (n = 5 for each sex) from the Mie region were used for the resequencing analysis and subjected to the PCR assay. (A) The results with the two sets of sex-identification primers for *H. discus hannai*. (B) The results with the two sets of sex-identification primers for *H. diversicolor*. M indicates the DNA size marker lane. These sex identification markers were validated in *H. discus hannai* and *H. diversicolor* from coastal areas of Japan (Figure S1, Table S1).

### 3.3. Sequencing and variant calling

The above results suggested that transferability of sex identification markers between species in the genus *Haliotis* is limited, prompting us to search for the sex-determining region in the genome of *H. gigantea.* We used a whole-genome resequencing strategy and obtained 99 Gb of clean reads for 20 individuals (n = 10 for each sex; Table S3). The reads were mapped to the draft genome of *H. gigantea*, with an average mapping rate of 97.7%. The average coverage for each sample was 3.98× (Table S3). Using ANGSD, a total of 40,353,502 SNPs were obtained. Since population sub-structure can bias the results of subsequent genetic analysis, we conducted a PCA of the SNP data. As expected for broodstock individuals obtained from the wild, we did not find substantial stratification associated with sex in the samples (Figure 3A).

**Figure 3.**
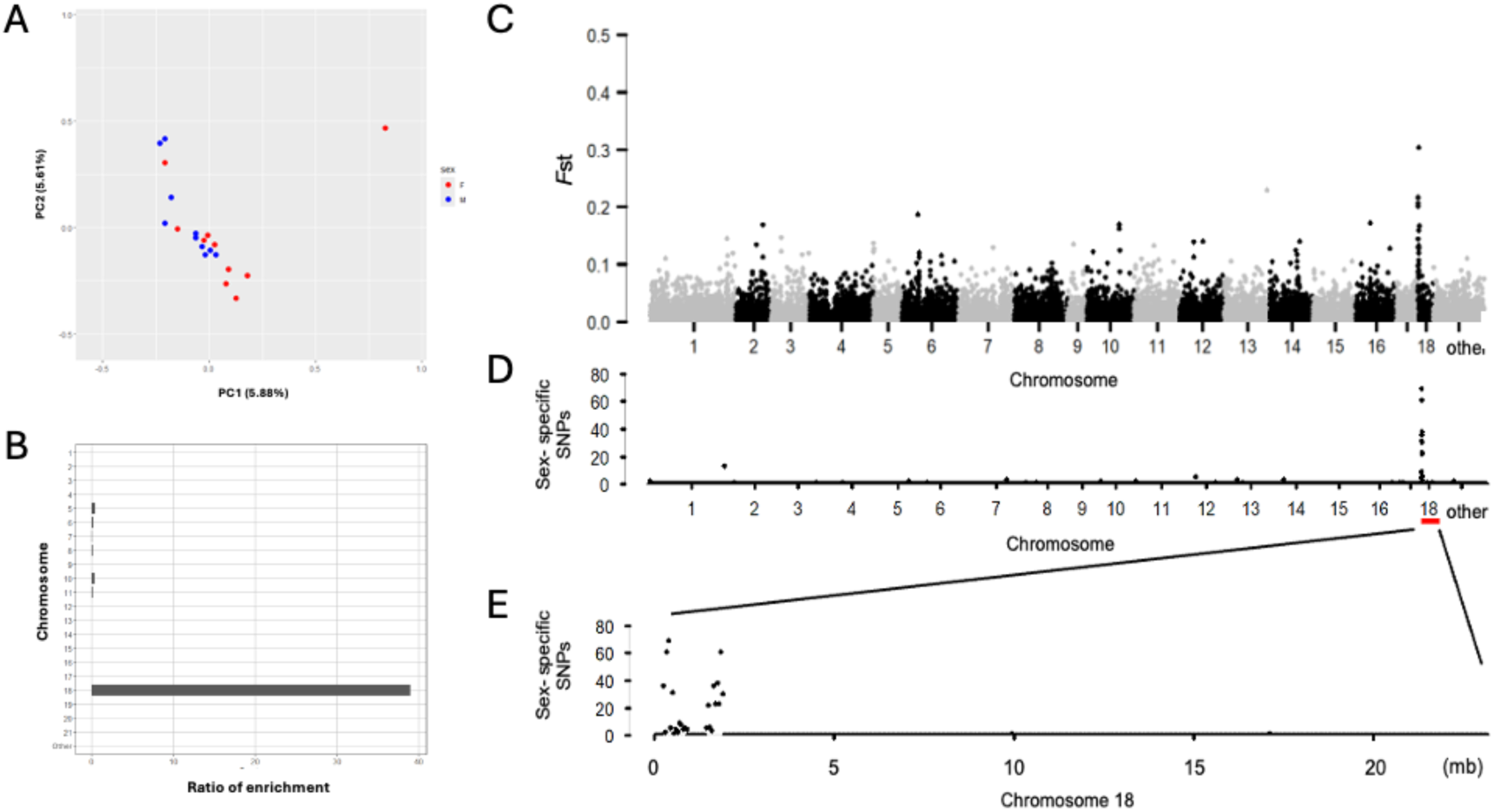
Genome-wide identification of the sex-determining locus in megai abalone (*H. gigantea*). (A) PCA plot showing no population structure associated with phenotypic sex in the tested abalones from the Mie region. Red circles indicate females and blue circles represent males. (B) The ratio of enrichment for sex-associated SNPs in each chromosome. SNPs exceeding the FDR threshold obtained from a genome-wide association study were counted. (C) *F*st analysis between males and females. (D) The number of sex-specific SNPs in a 50 kb window for all chromosomes. (E) The distribution of sex-specific SNPs along chromosome 18.

### 3.4. Identification of sex chromosome and sex determination locus

Sex chromosomes in fish can vary from being highly divergent (e.g., Chinese tongue sole) (Chen et al., 2014) to mostly identical (e.g., tiger pufferfish and greater amberjack) (Kamiya et al., 2012; Koyama et al., 2019). The degree of sequence divergence between sex chromosomes has a direct effect on the choice of approach for identifying the sex-determining region (Bewick et al., 2013; Palmer et al., 2019).

First, we performed GWAS using the 0.933 ANGSD package on 10 males and 10 females but failed to detect candidate sex chromosomes. The relatively small sample sizes may have reduced the power of the analysis. We then restricted the threshold for minor allele frequency from 0.2 to 0.3 to focus on SNP loci that are potentially fixed or nearly fixed on evolutionary Y (or W) chromosome. This analysis revealed that the number of associated SNPs exceeding the FDR threshold for phenotypic sex (3.113× 10^-4^) was more enriched on chromosome 18 than other chromosomes (Figure 3B).

The Fixation index (*F*st) is a powerful tool for detecting differentiation between diverging sex chromosomes using relatively small sample sizes (Gammerdinger et al., 2020; Rodrigues and Dufresnes, 2017). We therefore calculated the *F*st value in 50 kb windows between males and females in our sample. We found that chromosome 18 exhibited remarkable genetic differentiation between males and females (Figure 3C), suggesting that chromosome 18 is the sex chromosome of *H. gigantea*.

Diverging regions in young sex chromosome systems are expected to accumulate fixed sex-linked SNP loci that are heterozygous in one sex but homozygous in the other (i.e., sex-specific SNPs) (Bewick et al., 2013; Gammerdinger et al., 2016). We looked for such SNPs across all chromosomes and found 516 in total. Of these, 453 SNPs (87.8% of the total) were located on chromosome 18 (Figure 3D), and all of them on chromosome 18 were heterozygous in males but homozygous in females (Table 2). The result suggested that *H. gigantea* has an XX-XY sex chromosome system.

**Table 2.** The number and ratio of sex-specific SNPs in each chromosome in megai abalone (*H. gigantea*).

| Chromosome | Sex -<br>specific<br>SNPs | Male-specific<br>SNPs | Female-specific SNPs |
| --- | --- | --- | --- |
| Chr.1 | 2 (0.4%) | 1 (0.2%) | 1 (0.2%) |
| Chr.2 | 0 | 0 | 0 |
| Chr.3 | 2 (0.4%) | 2 (0.4%) | 0 |
| Chr.4 | 1 (0.2%) | 1 (0.2%) | 0 |
| Chr.5 | 2 (0.4%) | 2 (0.4%) | 0 |
| Chr.6 | 9 (1.7%) | 9 (1.7%) | 0 |
| Chr.7 | 3 (0.6%) | 3 (0.6%) | 0 |
| Chr.8 | 5 (1.0%) | 5 (1.0%) | 0 |
| Chr.9 | 2 (0.4%) | 1 (0.2%) | 1 (0.2%) |
| Chr.10 | 14 (2.7%) | 11 (2.1%) | 3 (0.6%) |
| Chr.11 | 1 (0.2%) | 0 | 1 (0.2%) |
| Chr.12 | 9 (1.7%) | 9 (1.7%) | 0 |
| Chr.13 | 2 (0.4%) | 1 (0.2%) | 1 (0.2%) |
| Chr.14 | 7 (1.4%) | 5 (1.0%) | 2 (0.4%) |
| Chr.15 | 0 | 0 | 0 |
| Chr.16 | 0 | 0 | 0 |
| Chr.17 | 0 | 0 | 0 |
| Chr.18 | 453 (87.8%) | 453 (87.8%) | 0 |
| Other | 4 (0.7%) | 0 | 4 (0.7%) |
| Total | 516 (100%) | 503 (97.5%) | 13 (2.5%) |

Furthermore, the density plot for the sex-specific SNPs along chromosome 18 revealed that the sex-determining region likely resides near one end of this chromosome (from the position 260,372 bp to 1,930,278 bp) (Figure 3E).

### 3.5. Development and verification of sex-specific markers

Indel polymorphisms are preferentially used as codominant genetic markers that can be easily detected by PCR and subsequent gel electrophoresis. We looked for sex-specific Indels in the diverged region on chromosome 18, which was characterized by the abundance of male-specific SNPs (the region from 260,372 bp to 1,930,278 bp)(Table S4). A 23-bp Indel was identified at position 280,120 bp. A visual inspection of the resequencing data using IGV confirmed that the 23-bp deletion was specific to males; therefore, this locus was heterozygous in all males but homozygous in all females (Figure 4). We designed PCRs to amplify the 440 bp region encompassing the 23-bp Indel locus.

**Figure 4.**
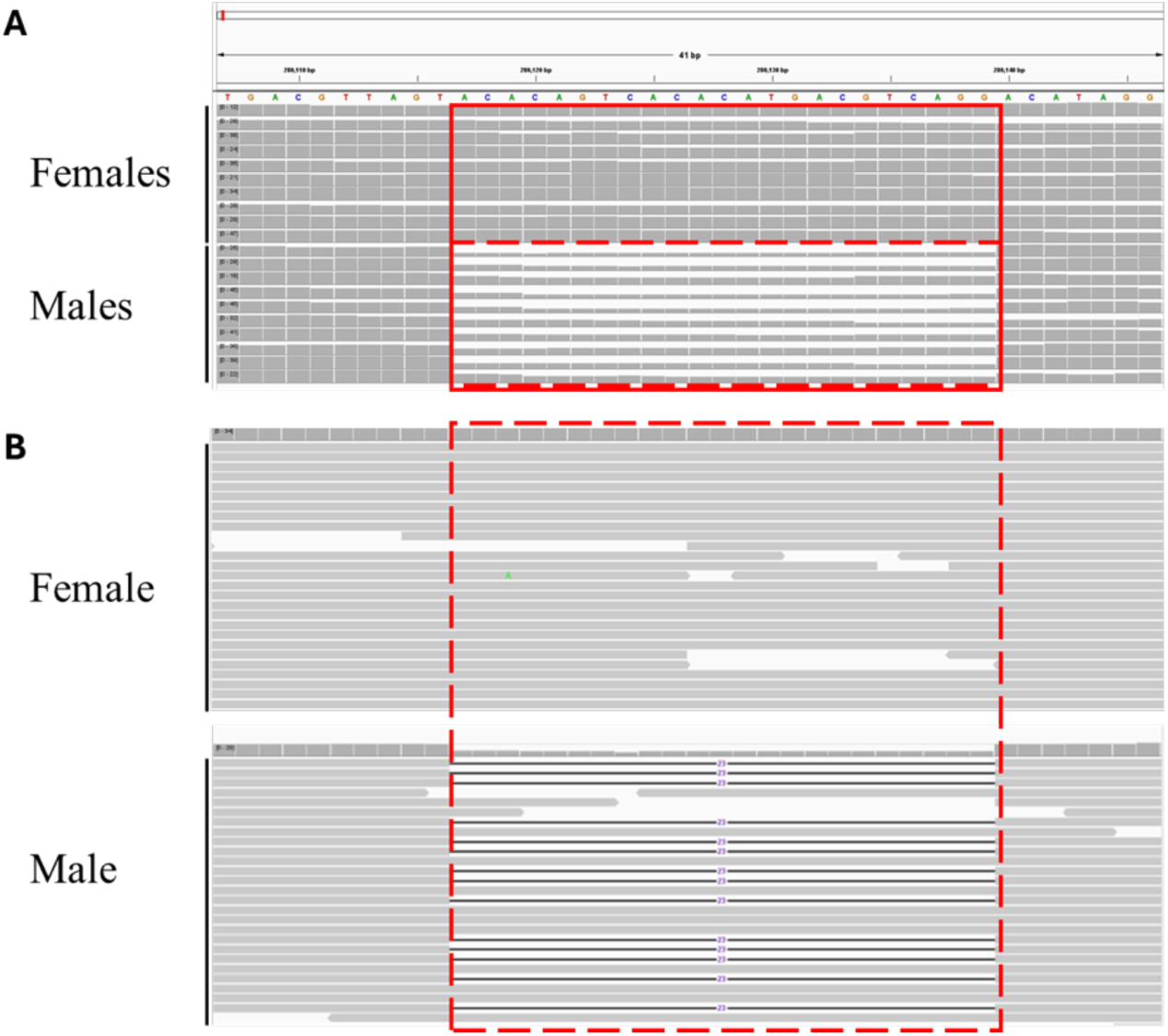
Visualization of alignment in the genomic region containing a male-specific deletion in megai abalone (*H. gigantea*). (A) Comparison of sequence depth among 20 individuals (10 females and 10 males). (B) Representative sequence alignments of a female and a male. The male-specific 23-base deletion is indicated as a horizontal black line with the number 23 (in purple).

We tested this candidate sex identification system using *H. gigantea* individuals from the Tokushima (n = 5 for each sex) and Mie (n = 5 for each sex) regions in addition to the Mie samples that were used for resequencing analysis (n = 5 for each sex). We found that PCR amplification resulted in two electrophoretic bands for phenotypic males but only one band for phenotypic females (Figure 5, see Figure S1C for resequenced samples). The genetic sex determined by the PCR analysis was consistent with the phenotypic observation of sex in both populations (Table 3) (Mie population *p* = 0.00001, Tokushima population *p* = 0.00793, Fisher’s exact tests).

**Figure 5.**
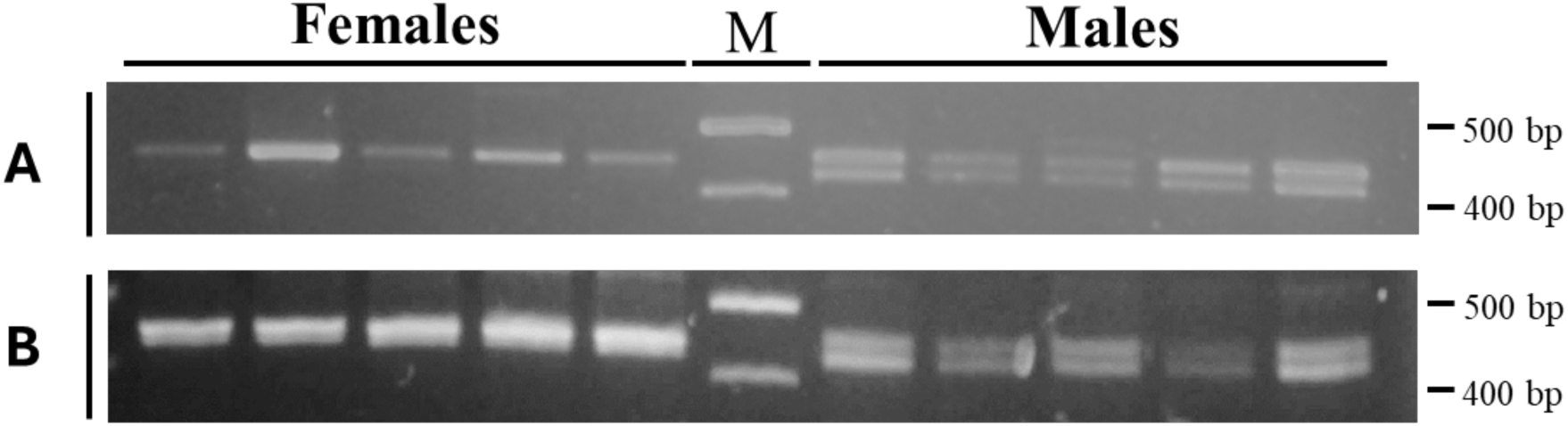
Validation of the sex identification maker in megai abalone (*H. gigantea*). (A) Five females and five males from the Mie region that had not been used for the resequencing analysis. (B) Five females and five males from the Tokushima region. M indicates the DNA size marker lane.

**Table 3.** Genotypes at the sex identification marker in megai abalone (*H. gigantea*) from two regions and their phenotypic sex.

|  | Tokushima population |  | Mie population |  |
| --- | --- | --- | --- | --- |
|  | Male | Female | Male | Female |
| **Heterozygous** | 5 | 0 | 10 | 0 |
| **Homozygous** | 0 | 5 | 0 | 10 |

### 3.6. Transferability of the diagnostic marker developed for *H. gigantea* to other abalone species

To test whether the sex identification marker for *H. gigantea* developed in this study could be used in *H. discus hannai* and *H. diversicolor*, we performed PCR on genomic DNA from both species (n = 5 for each sex, for each species). We detected no length polymorphisms in PCR products from *H. discus hannai*, although the PCR product was within the expected size range of 440 bp (Figure S1A). Moreover, no or unstable PCR amplification product was observed in *H. diversicolor* (Figure S1B). The results indicate that the male-specific deletion in *H. gigantea* identified in this study is unlikely to be shared by *H. discus hannai* and *H. diversicolor*.

## Discussion

In this study, we used a whole-genome resequencing approach to identify the sex chromosome and sex determination region of *H. gigantea* and this enabled the development of a genetic marker capable of distinguishing males and females. Our data suggested that males of this species are heterogametic and that there is a large region of sequence differentiation between the X and Y chromosomes. Furthermore, we showed that the sex identification markers so far identified in abalones are not transferable between species, highlighting the diversity of sex chromosomes within the genus *Haliotis*.

Previous cytological analyses showed that sex chromosomes vary from homomorphic to heteromorphic in animals (Becak et al., 1971; Devlin and Nagahama, 2002; Ohno, 1967). Even within seemingly homomorphic sex chromosome systems, recent studies have identified sequence diversification between X and Y or Z and W chromosomes (Palmer et al., 2019). Although previous studies suggested the absence of heteromorphic sex chromosomes in *H. gigantea* (Miyaki et al., 1997), as in the case of other abalones (Escárate & Portilla, 2007; Gallardo et al., 2005; López et al., 2017; Okumura et al., 1999), our analysis here indicated that the sex chromosomes of *H. gigantea* have a diverged region spanning at least 2 Mb (Figure 3E). The distribution of fixed SNP alleles in this region of the Y chromosome (Figure 3E) indicates that there is no recombination between the X and Y chromosomes; however, large-scale sequence degeneration has not yet occurred since short reads could be aligned to the draft genome sequence without prominent differences in coverage between sexes. The female/male ratio of the read coverage was 1.10 (SD 0.27) and 1.05 (SD 0.19) in the diverged region (260372 bp ‒ 1930278 bp) and the pseudoautosomal regions, respectively (Figure S2). In the diverged region, our search for Indel polymorphisms identified only one locus consistently associated with sex (Table S4). The low frequency of Indel loci is likely due to the low coverage of the resequencing data and misalignments around Indels. Therefore, a larger number of male-specific Indel loci might be recovered using more extensive data. Nevertheless, we successfully developed a sex identification marker that could be applied to geographically distant populations (Table 3). It is notable that the cytologically homomorphic sex chromosomes in both *H. discus hannai* and *H. diversicolor* (Arai et al., 1988; Okumura et al., 1999) have recently been shown to have considerable sequence differences (Luo et al., 2021; Weng et al., 2022).

Although sex identification markers have been developed for many economically important fish, some of these are specific to a few families or stock populations (Agawa et al., 2015; Eisbrenner et al., 2014; Taslima et al., 2020), probably due to population structure, kinship, or introgression. A previous study based on genome-wide SNP data for *H. gigantea* indicated that there is no clear genetic differentiation among three groups from geographically distant sites along the coast of the Japanese archipelago (Hirase et al., 2021). The absence of genetic structure is likely due to wide dispersion during the planktonic larval stage (Miyake et al., 2009), as in other abalone species (Chambers et al., 2006; Miyake et al., 2017). Therefore, the sex-determining region on chromosome 18 may be shared in most of wild populations of *H. gigantea* in Japan and hence our sex identification marker will widely applicable. However, some caution is necessary as sex-determining loci can evolve in a genetically isolated population in a relatively short period of time (Nanda et al., 2003; Shinomiya et al., 2010; Wilson et al., 2014; Martínez et al., 2021). Hybridization between closely-related species or diverged populations can also result in the emergence of new sex-determining regions (Dixon et al., 2019; Franchini et al., 2018; Kabir et al., 2022).

Moreover, given the high genetic diversity of the genus *Haliotis* that has been revealed by genomic analyses (Hirase et al., 2021; Hirase et al., 2023; Nam et al., 2021; Wooldridge et al., 2024), the Indel variants we identified may not be present in some populations. Further studies using genome resequencing data from diverse populations will be necessary to determine whether this is the case.

It has been shown that sex chromosomes have evolved independently on a number of occasions in animal species; moreover, this type of evolution has occurred even within closely related species in some finfish lineages (e.g., Kabir et al., 2022; Myosho et al., 2015; Ross et al., 2009). A recent study in molluscs identified the independent evolution of sex chromosomes among distantly related lineages of scallops (family *Pectinidae*) (Han et al., 2022). For abalones (family *Haliotidae*), Weng et al., (2022) discussed the possibility that the sex chromosomes are not conserved between the distantly related species *H. discus hannai* and *H. diversicolor*, which belong to two deeply divergent clades: the North Pacific clade and the European-Australasian clade, respectively (Geiger, 2000; Geiger and Thacker, 2005; Streit et al., 2005). Our PCR analysis revealed that the sex identification markers are not reciprocally transferable even between the closely related species *H. discus hannai* and *H. gigantea* (Figure 1 and Figure S1). This non-reciprocity occurred despite their similarities in genomic sequences (Hirase et al., 2021) and, hence, likely conservation of primer loci sequences. Indeed, a blast search of *H. discus hannai* primers against the *H. gigantea* draft genome indicated that the loci corresponding to the four primers for sex chromosome markers are located on an autosome (chromosome 14) of *H. gigantea* with 100% match (Table S5). Therefore, the two closely related species likely have different sex determination regions, suggesting a rapid turnover of sex chromosomes or sex-determining regions has occurred in abalones as in some lineages of finfish such as medaka and its relatives (Myosho et al., 2015). However, our analyses were limited by insufficient resources for comparative genomics. Currently, a fragmented genome assembly of *H. discus hannai* has been deposited in the NCBI database (scaffold N50 = 211,346 bp) (Nam et al., 2017). However, no chromosome-level genome assemblies are publicly available for any of the western Pacific abalones or *H. diversicolor*. Elucidation of the nature of the turnover of sex chromosomes among these abalone species will require the availability of contiguous genome assemblies.

## Acknowledgments

We are grateful to the laboratory members for their valuable feedback on this research, particularly to S. Hosoya, A. Kabir, and S. Kato for their assistance with genetic analyses. We thank N. Mizuno for setting up a breeding tank.

## Funding

This work was supported by the Grants-in-Aid for Scientific Research (grant numbers 22H00377 and 23K05267).

---

**Figure S1.**
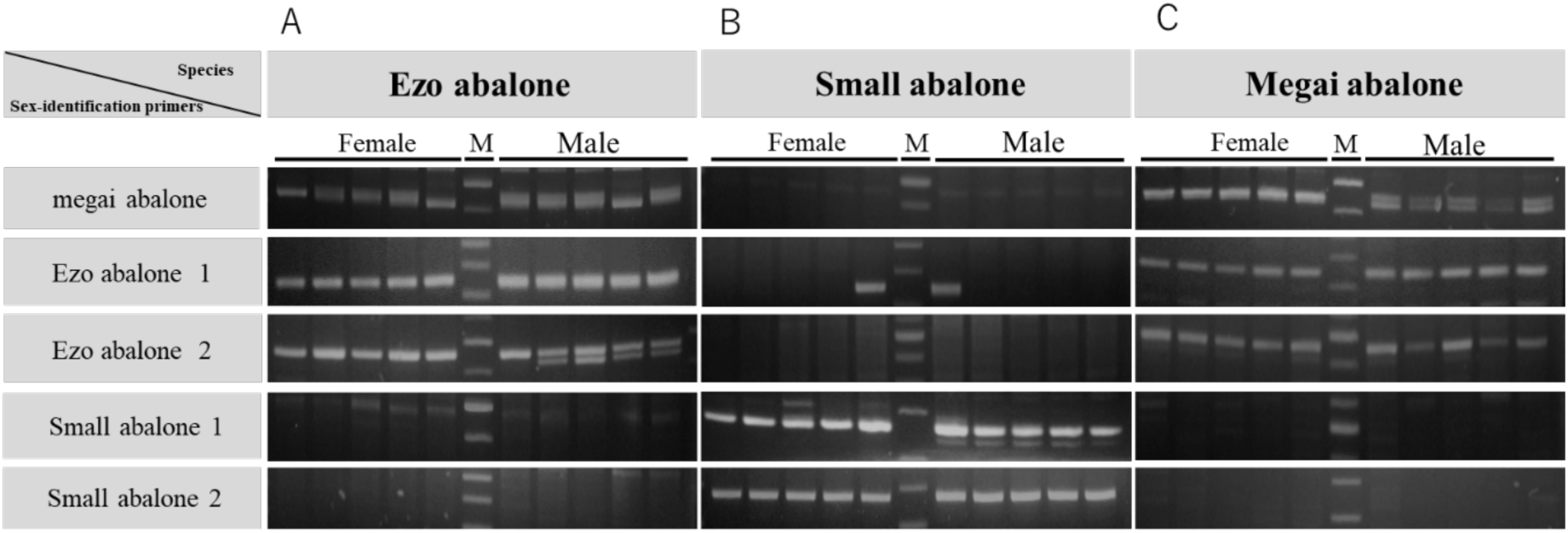
PCR products from Ezo abalone (*Haliotis discus hannai*), small abalone (*H. diversicolor*), and megai abalone (*H. gigantea*) with the sex-identification primers for the three species. (A) *H. discus hannai* samples. (B) *H. diversicolor* samples. (C) *H. gigantea* samples from Mie used for resequencing analysis. M indicates the DNA size marker lane. The sex-identification primers for Ezo abalone 1 and 2, respectively, correspond to MSP2 and MSP7 reported in Luo et al. (2021). The sex-identification primers for small abalone 1 and 2, respectively, correspond to SC9 and SD3 reported in Weng et al. (2022).

**Figure S2.**
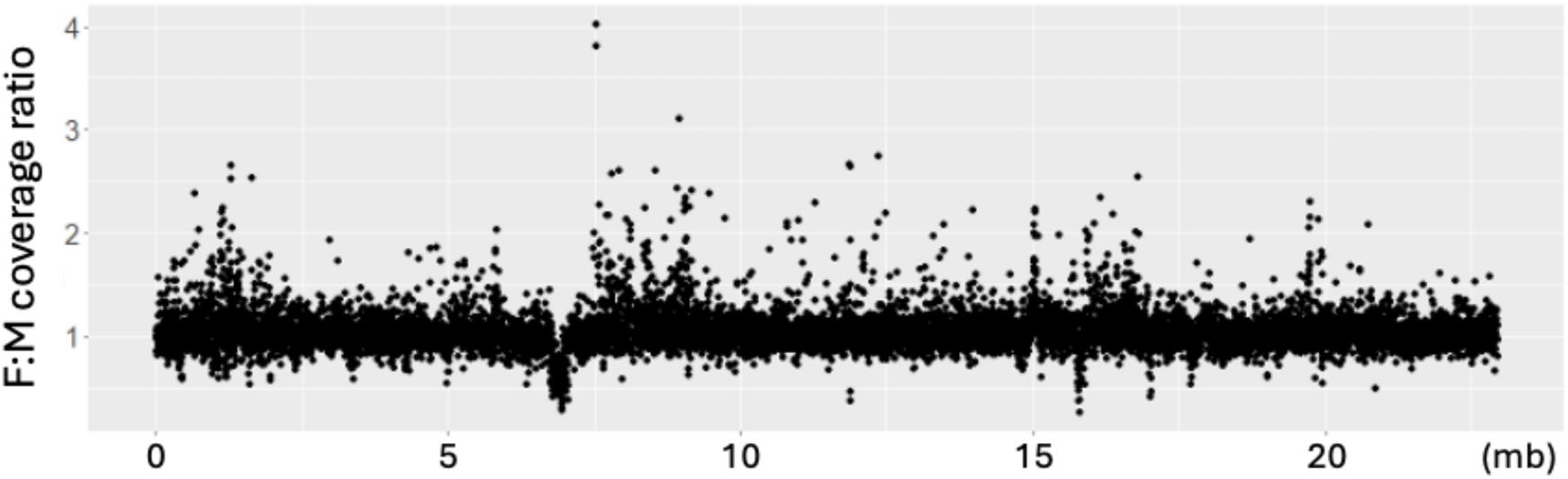
The female/male ratio of DNAseq read coverage along the sex chromosome in megai abalone (*H. gigantea*). Resequencing reads from 10 females and 10 males were aligned against the draft genome sequence. The relative depth of coverage between sexes at a window size of 2 kb was plotted.

**Table S1.** Sample information of *Haliotis gigantea* , *H. discus hannai* and *H. diversicolor* used in this study.

| Sample ID | Species | Sampling region | Sample type | Shell length | Phenotypic sexes | Experimental use |
| --- | --- | --- | --- | --- | --- | --- |
| Mie_1 | *H. gigantea* | Mie prefecture in Japan | Broodstock | About 100.0mm | Female | Resequencing |
| Mie_2 | *H. gigantea* | Mie prefecture in Japan | Broodstock | About 100.0mm | Female | Resequencing |
| Mie_3 | *H. gigantea* | Mie prefecture in Japan | Broodstock | About 100.0mm | Female | Resequencing |
| Mie_4 | *H. gigantea* | Mie prefecture in Japan | Broodstock | About 100.0mm | Female | Resequencing |
| Mie_5 | *H. gigantea* | Mie prefecture in Japan | Broodstock | About 100.0mm | Female | Resequencing |
| Mie_6 | *H. gigantea* | Mie prefecture in Japan | Broodstock | About 100.0mm | Female | Resequencing |
| Mie_7 | *H. gigantea* | Mie prefecture in Japan | Broodstock | About 100.0mm | Female | Resequencing |
| Mie_8 | *H. gigantea* | Mie prefecture in Japan | Broodstock | About 100.0mm | Female | Resequencing |
| Mie_9 | *H. gigantea* | Mie prefecture in Japan | Broodstock | About 100.0mm | Female | Resequencing |
| Mie_10 | *H. gigantea* | Mie prefecture in Japan | Broodstock | About 100.0mm | Female | Resequencing |
| Mie_11 | *H. gigantea* | Mie prefecture in Japan | Broodstock | About 100.0mm | Male | Resequencing |
| Mie_12 | *H. gigantea* | Mie prefecture in Japan | Broodstock | About 100.0mm | Male | Resequencing |
| Mie_13 | *H. gigantea* | Mie prefecture in Japan | Broodstock | About 100.0mm | Male | Resequencing |
| Mie_14 | *H. gigantea* | Mie prefecture in Japan | Broodstock | About 100.0mm | Male | Resequencing |
| Mie_15 | *H. gigantea* | Mie prefecture in Japan | Broodstock | About 100.0mm | Male | Resequencing |
| Mie_16 | *H. gigantea* | Mie prefecture in Japan | Broodstock | About 100.0mm | Male | Resequencing |
| Mie_17 | *H. gigantea* | Mie prefecture in Japan | Broodstock | About 100.0mm | Male | Resequencing |
| Mie_18 | *H. gigantea* | Mie prefecture in Japan | Broodstock | About 100.0mm | Male | Resequencing |
| Mie_19 | *H. gigantea* | Mie prefecture in Japan | Broodstock | About 100.0mm | Male | Resequencing |
| Mie_20 | *H. gigantea* | Mie prefecture in Japan | Broodstock | About 100.0mm | Male | Resequencing |
| Mie_21 | *H. gigantea* | Mie prefecture in Japan | Broodstock | About 100.0mm | Female | PCR for genetic sex determination |
| Mie_22 | *H. gigantea* | Mie prefecture in Japan | Broodstock | About 100.0mm | Female | PCR for genetic sex determination |
| Mie_23 | *H. gigantea* | Mie prefecture in Japan | Broodstock | About 100.0mm | Female | PCR for genetic sex determination |
| Mie_24 | *H. gigantea* | Mie prefecture in Japan | Broodstock | About 100.0mm | Female | PCR for genetic sex determination |
| Mie_25 | *H. gigantea* | Mie prefecture in Japan | Broodstock | About 100.0mm | Female | PCR for genetic sex determination |
| Mie_26 | *H. gigantea* | Mie prefecture in Japan | Broodstock | About 100.0mm | Male | PCR for genetic sex determination |
| Mie_27 | *H. gigantea* | Mie prefecture in Japan | Broodstock | About 100.0mm | Male | PCR for genetic sex determination |
| Mie_28 | *H. gigantea* | Mie prefecture in Japan | Broodstock | About 100.0mm | Male | PCR for genetic sex determination |
| Mie_29 | *H. gigantea* | Mie prefecture in Japan | Broodstock | About 100.0mm | Male | PCR for genetic sex determination |
| Mie_30 | *H. gigantea* | Mie prefecture in Japan | Broodstock | About 100.0mm | Male | PCR for genetic sex determination |
| Tokushima_1 | *H. gigantea* | Tokushima prefecture in Japan | Wild | 107.7mm | Female | PCR for genetic sex determination |
| Tokushima_2 | *H. gigantea* | Tokushima prefecture in Japan | Wild | 108.9mm | Female | PCR for genetic sex determination |
| Tokushima_3 | *H. gigantea* | Tokushima prefecture in Japan | Wild | 109.5mm | Female | PCR for genetic sex determination |
| Tokushima_4 | *H. gigantea* | Tokushima prefecture in Japan | Wild | 115.6mm | Female | PCR for genetic sex determination |
| Tokushima_5 | *H. gigantea* | Tokushima prefecture in Japan | Wild | 123.8mm | Female | PCR for genetic sex determination |
| Tokushima_6 | *H. gigantea* | Tokushima prefecture in Japan | Wild | 106.8mm | Male | PCR for genetic sex determination |
| Tokushima_7 | *H. gigantea* | Tokushima prefecture in Japan | Wild | 110.4mm | Male | PCR for genetic sex determination |
| Tokushima_8 | *H. gigantea* | Tokushima prefecture in Japan | Wild | 105.6mm | Male | PCR for genetic sex determination |
| Tokushima_9 | *H. gigantea* | Tokushima prefecture in Japan | Wild | 111.9mm | Male | PCR for genetic sex determination |
| Tokushima_10 | *H. gigantea* | Tokushima prefecture in Japan | Wild | 106.3mm | Male | PCR for genetic sex determination |
| Ezo_1 | *H. discus hannai* | Hokkaido in Japan | Wild | 78.6mm | Female | PCR for genetic sex determination |
| Ezo_2 | *H. discus hannai* | Hokkaido in Japan | Wild | 90.1mm | Female | PCR for genetic sex determination |
| Ezo_3 | *H. discus hannai* | Hokkaido in Japan | Wild | 85.9mm | Female | PCR for genetic sex determination |
| Ezo_4 | *H. discus hannai* | Hokkaido in Japan | Wild | 87.9mm | Female | PCR for genetic sex determination |
| Ezo_5 | *H. discus hannai* | Hokkaido in Japan | Wild | 77.7mm | Female | PCR for genetic sex determination |
| Ezo_6 | *H. discus hannai* | Hokkaido in Japan | Wild | 85.1mm | Male | PCR for genetic sex determination |
| Ezo_7 | *H. discus hannai* | Hokkaido in Japan | Wild | 85.8mm | Male | PCR for genetic sex determination |
| Ezo_8 | *H. discus hannai* | Hokkaido in Japan | Wild | 84.4mm | Male | PCR for genetic sex determination |
| Ezo_9 | *H. discus hannai* | Hokkaido in Japan | Wild | 86.1mm | Male | PCR for genetic sex determination |
| Ezo_10 | *H. discus hannai* | Hokkaido in Japan | Wild | 79.0mm | Male | PCR for genetic sex determination |
| Small_1 | *H. diversicolor* | Oita prefecture in Japan | Wild | 58.2mm | Female | PCR for genetic sex determination |
| Small_2 | *H. diversicolor* | Oita prefecture in Japan | Wild | 62.8mm | Female | PCR for genetic sex determination |
| Small_3 | *H. diversicolor* | Oita prefecture in Japan | Wild | 54.4mm | Female | PCR for genetic sex determination |
| Small_4 | *H. diversicolor* | Oita prefecture in Japan | Wild | 59.7mm | Female | PCR for genetic sex determination |
| Small_5 | *H. diversicolor* | Oita prefecture in Japan | Wild | 56.7mm | Female | PCR for genetic sex determination |
| Small_6 | *H. diversicolor* | Oita prefecture in Japan | Wild | 69.5mm | Male | PCR for genetic sex determination |
| Small_7 | *H. diversicolor* | Oita prefecture in Japan | Wild | 61.0mm | Male | PCR for genetic sex determination |
| Small_8 | *H. diversicolor* | Oita prefecture in Japan | Wild | 67.9mm | Male | PCR for genetic sex determination |
| Small_9 | *H. diversicolor* | Oita prefecture in Japan | Wild | 66.0mm | Male | PCR for genetic sex determination |
| Small_10 | *H. diversicolor* | Oita prefecture in Japan | Wild | 70.4mm | Male | PCR for genetic sex determination |

**Table S2.**
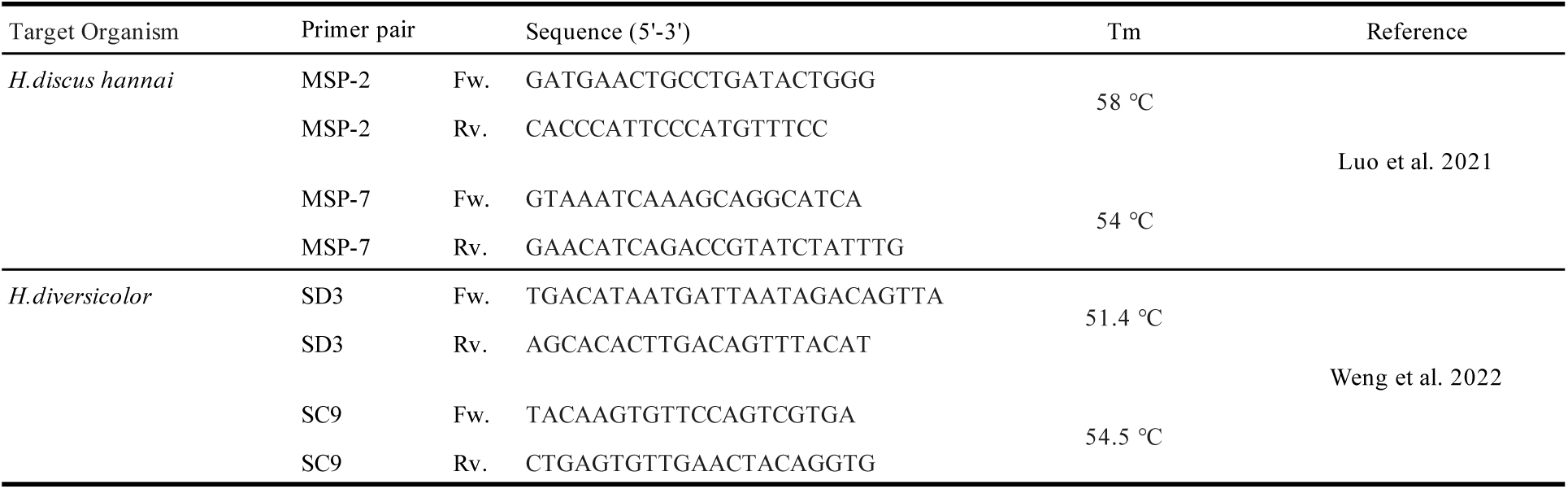
Sex-identification primers for Ezo abalone (*Haliotis dis cus hannai* ) and small abalone (*H. diversicolor* ) developed in previous reports.

**Table S3.** Genome resequencing statistics of *Haliotis gigantea*.

| ID | Phenotypic Sex | Clean Read | Mapping Percentage (%) | Coverage (X) |
| --- | --- | --- | --- | --- |
| Mie_1 | Female | 48,804,168 | 97.60 | 1.8 |
| Mie_2 | Female | 72,565,973 | 96.50 | 3.1 |
| Mie_3 | Female | 109,577,379 | 97.71 | 4.9 |
| Mie_4 | Female | 72,319,516 | 97.96 | 3.1 |
| Mie_5 | Female | 148,863,656 | 97.67 | 7.0 |
| Mie_6 | Female | 87,682,829 | 97.90 | 3.6 |
| Mie_7 | Female | 110,060,936 | 98.32 | 3.2 |
| Mie_8 | Female | 111,176,475 | 98.42 | 3.4 |
| Mie_9 | Female | 122,340,909 | 97.76 | 5.3 |
| Mie_10 | Female | 124,252,423 | 97.93 | 5.1 |
| Mie_11 | Male | 93,448,424 | 97.68 | 3.1 |
| Mie_12 | Male | 108,544,893 | 98.27 | 3.8 |
| Mie_13 | Male | 116,802,218 | 98.30 | 4.2 |
| Mie_14 | Male | 83,546,687 | 94.92 | 3.1 |
| Mie_15 | Male | 102,577,478 | 97.95 | 4.6 |
| Mie_16 | Male | 94,943,341 | 97.81 | 4.6 |
| Mie_17 | Male | 98,607,557 | 97.95 | 4.0 |
| Mie_18 | Male | 83,911,223 | 98.20 | 3.2 |
| Mie_19 | Male | 104,521,745 | 97.98 | 4.1 |
| Mie_20 | Male | 98,718,580 | 98.07 | 4.3 |

**Table S4.** The number of INDELs in each chromosome in megai abalone (H. gigantea).

| **Chr** | **INDELs** | **Male-specific INDELs** |
| --- | --- | --- |
| Chr.1 | 2941 (4.6%) | 0 |
| Chr.2 | 5080 (7.9%) | 0 |
| Chr.3 | 1670 (2.6%) | 0 |
| Chr.4 | 2164 (3.4%) | 0 |
| Chr.5 | 2298 (3.6%) | 0 |
| Chr.6 | 3920 (6.1%) | 0 |
| Chr.7 | 3782 (5.9%) | 0 |
| Chr.8 | 3374 (5.2%) | 0 |
| Chr.9 | 1797 (2.8%) | 0 |
| Chr.10 | 6457 (10.0%) | 0 |
| Chr.11 | 3658 (5.7%) | 0 |
| Chr.12 | 3594 (5.6 %) | 0 |
| Chr.13 | 3793 (5.9%) | 0 |
| Chr.14 | 3924 (6.1%) | 0 |
| Chr.15 | 3170 (4.9%) | 0 |
| Chr.16 | 4363 (6.8%) | 0 |
| Chr.17 | 3676 (5.7 %) | 0 |
| Chr.18 | 1371 (2.1%) | 1 |
| Other | 3483 (5.4%) | 0 |
| Total | 64515 (100%) | 1 |

**Table S5.** Blast reports of the sex-identification primers for *Haliotis discus hannai* against the draft genome sequence of *H. gigantea*.

| Sex-identification primers for *H.discus.hannai* |  |  | Genomic positions in the draft genome of *H. gigantea* |  |  |  |  |  |  |
| --- | --- | --- | --- | --- | --- | --- | --- | --- | --- |
| Primer name | Start | End | Chromosome | Start | End | Strand | E-value | Sequence identity (%) | Expected length (bp) of the PCR product |
| MSP2_Fw | 1 | 21 | 14 | 1,723,957 | 1,723,937 | minus | 0.004 | 100 | 251 |
| MSP2_Rv | 1 | 19 | 14 | 1,723,724 | 1,723,706 | minus | 0.054 | 100 |  |
| MSP7_Fw | 1 | 20 | 14 | 539,028 | 539,047 | plus | 0.015 | 100 | 467 |
| MSP7_Rv | 1 | 23 | 14 | 539,473 | 539,495 | plus | 0.0003 | 100 |  |

## Notes

### Competing Interest Statement

The authors have declared no competing interest.

